# Cross—reactivity of antibodies against microbial proteins to human tissues as basis of Crohn’s disease and other autoimmune diseases

**DOI:** 10.1101/116574

**Authors:** Peilin Zhang, Lawrence M. Minardi, J. Todd Kuenstner, Stephen M. Zekan, Feng Zhu, Yinling Hu, Rusty Kruzelock

## Abstract

**Background:** Autoimmune disease is generally a systemic inflammatory response with production of autoantibodies. In this study, we investigated the anti-microbial antibodies in circulation in cases of Crohn’s disease (CD), Sjogren’s syndrome (SS) and other autoimmune disease and their roles in the pathogenesis of these autoimmune diseases.

**Material and methods:** Western blot was used to determine the reactivity of human plasmas from patients with CD and SS as the primary antibodies against the whole microbial extracts. The microbial proteins reactive to patients’ plasma were further identified and the modified sandwich ELISA assays were used to determine the blood levels of antibodies against these microbial proteins in patients with CD and SS. Antibodies against the microbial proteins are used for immunohistochemical staining of normal human tissue.

**Results:** A group of 7 microbial proteins was identified reactive to the plasmas of patients with CD and SS including DNA-directed RNA polymerase B (RPOB), and elongation factor G (EF-G) from *Staphylococcus aureus* and *Staphylococcus pseudintermedius* (*S. aureus* and *S. pseudintermedius*), ATP synthase alpha (ATP5a) and heat shock protein 65 (*Hsp65*) from *Mycobacterium avium subspecies paratuberculosis* (MAP), elongation factor Tu (EF-Tu) and outer membrane porin C (ompC) from *Escherichia Coli* (*E. coli*). Anti-microbial antibodies can cross-react to normal human tissues. The levels of antibodies against the microbial proteins are significantly elevated in the patients with CD and SS.

**Conclusion:** The levels of antibodies against the microbial proteins are significantly elevated in CD and SS. The cross-reactivity of the anti-microbial antibodies to human tissue provides a new mechanism of pathogenesis of autoimmune diseases such as CD and SS.

---

Autoimmune diseases are a spectrum of clinical conditions characterized by systemic manifestation with circulating antibodies against the patient’s own tissues with the exception of Crohn’s disease in which no identifiable autoantibody is known(1). Genetic susceptibility is an essential pre-requisite condition for the pathogenesis of autoimmune diseases, although the molecular mechanisms of most autoimmune diseases are unclear (1). Recent advances of microbiome research demonstrated that the surface area of human body including body cavities is covered by a plethora of microbes (commensal), and these microbes are essential in host defense against pathogens, innate and adaptive immunity development and prevention of a variety of diseases (2, 3). The symbiotic relationship of the human body with its own microbes becomes an interesting issue in the pathogenesis of infection, immunity and various degenerative diseases.

Decades ago the discovery of anti-streptolysin O antibody in the circulation of patients with streptococcal infection (pharyngitis) cross reacting with human valvular tissue and renal glomerular tissue leading to valvular heart disease and glomerulonephritis provides the basis of understanding the mechanism of streptococcal infection and the related disease process, which led to new treatment and preventive strategy (4). Immunological cross reactivity between the Streptococcal M protein and cardiac myosin/tropomyosin induces myocardial damage with reactive T- and B-lymphocytes, and there are shared epitopes between the M protein and the cardiac myosin (5–7). A similar mechanism has been described for Epstein-Barr viral infection and hemolytic anemia (8). In our effort to determine the role of mycobacterial infection in Crohn’s disease, we found there are elevated antibody levels in the patient’s circulation against a panel of microbial proteins. These microbes are commonly present on the body surface of human or in the environment. These anti-microbial antibodies can cross-react with human tissue, potentially leading to dysfunction of human tissue and various clinical conditions. The elevated levels of anti-microbial antibodies can be used as diagnostic tools for assessment of the relationship and compatibility between the host and its own microbes, and the presence of anti-microbial antibodies provide a new direction of research in understanding the mechanism of autoimmune diseases.

## Material and Method

### 1 Western blot analysis using the microbial cell extracts and the patients’ plasmas as primary antibodies

In order to determine if the human plasma from the patients with Crohn’s disease and Sjogren’s syndrome react to the microbial proteins, the whole microbial extracts from *Staphylococcus aureus, Staphylococcus pseudintermedius, Escherichia coli, Mycobacterium avium subspecies paratuberculosis (MAP)* and *Mycobacterium avium subspecies hominissuis (MAH)* were prepared and Western blot analysis was performed as described (9–11). A strain of *S. aureus* was isolated from our own lab using high osmolarity culture medium based on Middlebrook 7H9 broth (BD Difco Cat. # DF0713-17-9) from a patient with history of methicillin-resistant *S. aureus* infection, and this strain of *S. aureus* was maintained in Tryptic Soy broth/plate (Sigma Aldrich Cat. # 22092). The isolate had been identified by 16S PCR/sequencing as previously described (9–11), and the subsequently by whole genomic sequencing using Illumina Miseq service from West Virginia University Genomic Core Facility (data not shown). The strain of *S. pseudintermedius* was isolated from a skin wound of a domestic dog with moist dermatitis using the Tryptic Soy broth and the identity of the bacteria was confirmed by 16S PCR/sequencing (9–11) and subsequently whole genomic sequencing at WVU Genomic Core Facility (Data not shown). MAP strain (MAP *Dominic* was obtained commercially from ATCC (Cat: 43545). MAH strain was isolated from a patient with Crohn’s disease and this MAH strain was characterized previously(9–11), and the identity was confirmed by whole genomic sequencing at WVU Genomic Core Facility. *S. aureus* and *S. pseudintermedius* were maintained in the liquid Tryptic Soy broth and MAP and MAH were maintained in the Middlebrook 7H9 liquid medium with supplement of mycobactin J (2 ug/ml) as described (10, 11). A strain of *Escherichia coli* was from ATCC (Cat. # 25922) and maintained on the L-Broth (BD Difco Cat. # DF0478-17-4). The bacterial protein extracts were prepared using the Quick & Easy Protein Extraction kit by PZM Diagnostic (Cat. # QEPEKS, Charleston, WV) according to the manufacturer’s instruction. Briefly, all microbial cultures (1.0 ml) in the appropriate media were transferred to a 1.5 ml microcentrifuge tubes, and the cultures were centrifuged at 15,000 g for 5 minutes. The supernatants were discarded, and the microbial cell pellets were washed with phosphate-buffered saline (PBS) once. The microbial cell pellets were washed with Solution #1 and #2 once respectively, and the cell pellets were resuspended in Solution #3 containing 1% SDS at 4°C overnight to extract cellular proteins. The protein concentration is determined by using Qubit protein quantification kit (Fisher Scientific). The whole microbial extracts (20 ug) were subjected to 10% SDS-polyacrylamide gel electrophoresis with molecular weight marker, and the cellular proteins were electroblotted onto nitrocellulose membrane. The microbial proteins on the nitrocellulose membrane were incubated with the plasmas (1 to 50 dilution) from patients with Crohn’s disease at room temperature for 2 hours (or 4°C overnight), and secondary antibodies and detection systems were provided by using the Pierce enhanced chemiluminescent detection kit (Fisher Scientific) according to the manufacturer’s instruction.

### 2 Identification of the microbial proteins by immunoprecipitation and mass spectrometry

Seven microbial proteins showed in Figure 1 were precipitated by using the same patient’s plasmas as the primary antibodies. Briefly, the bacterial/mycobacterial cell extracts (50 ug/200ul) were incubated with 50 ul plasma from the patients and 50 ul protein A-coupled agarose beads (Pierce Corp, Thermo Fisher) at 4°C overnight in the 1.5 ml microcentrifuge tubes. The reaction tubes were briefly centrifuged at 6000 g for 1 minute, and the supernatants were carefully removed, and the agarose beads were washed 6 times with phosphate buffered saline with 0.2% Tween 20 (PBST). The specific microbial proteins binding to the plasmas were captured by using the protein A-agarose column (Pierce from Thermo Fisher) and the captured microbial proteins were analyzed by Nanospray liquid chromatography/mass spectrometry analysis (LC/MS/MS) with database search by Poochon Scientific, LLC, Frederick, MD through contract work. A list of microbial proteins was obtained from each immunoprecipitation tube, and only the proteins of similar molecular weights were of interest for further validation test (Table 1).

**Figure 1:**
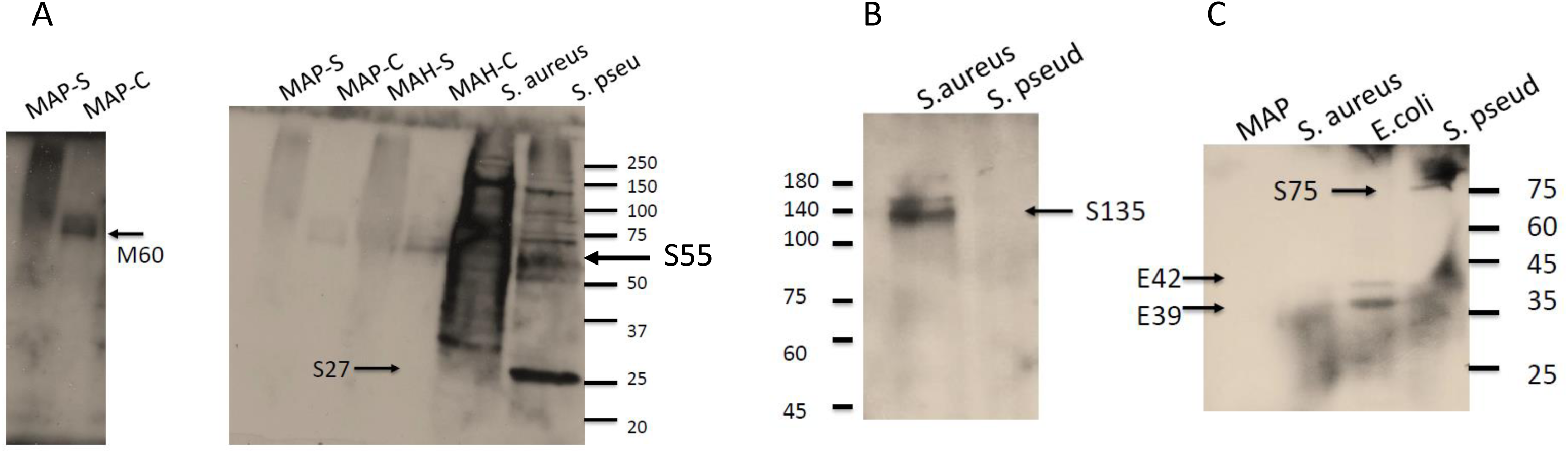
Identification of microbial proteins using Western blot analysis and the plasma of patients with Crohn’s disease. Various microbial cell extracts were separated by SDS-polyacrylamide gel electrophoresis, and transferred onto the nitrocellulose membrane. The protein extracts on the membranes were incubated with the plasma (1 to 50 dilutions) from the patients with Crohn’s disease. Three patients’ plasmas, A, B and C were used for analyses. The signals were detected with the secondary antibody against human IgG with HRP conjugates and enhanced chemiluminescent detection system (ECL). Panel A, B and C were using the plasmas from patients A, B and C. MAP-S, MAP culture supernatant, MAP-C, MAP culture cell pellet, MAH-S, MAH culture supernatant, MAH-C, MAH culture cell pellet, *S pseud-S pseudintermedius*.

**Table 1:** List of the microbial proteins is identified by Western blot analysis, immunoprecipitation and mass spectrometry. The same patient’s plasma was used as for the Western blot analysis in Figure 1 for immunoprecipitation. The precipitated microbial proteins are analyzed by mass spectrometry, and the database search matched the list of proteins by both peptide identities and molecular weight. Mass spectrometry study was performed by Poochon Scientific LLC, Frederick, MD. Genbank accession numbers were followed after the proteins names. Homology to human homologues was by BLASTP search at https://blast.ncbi.nlm.nih.gov/Blast.cgi

| • ID | Full name | Homology to human (By BLASTP) |
| --- | --- | --- |
| • S135 | DNA-directed RNA polymerase subunit B from *S. aureus* (RPOB)(#ANR83811) | 47% |
| • S75 | Elongation factor G from *S. pseudintemedius* (EF-G) (# WP_014614745) | 61% |
| • M60 | Heat shock protein 65 from *M. paratuberculosis* (HSP65) (# P9WPE6) | 68% |
| • S55 | ATP synthase subunit alpha from *S. pseudintermedius* (ATP5a)(#KZK18041) | 74% |
| • E42 | Elongation factor Tu from *Escherichia Coli* (EF-Tu) (# AAA50993) | 75% |
| • E39 | Outer membrane porin protein C from *Escherichia Coli* (ompC) (#AAA23728) | No human homologue |
| • S27 | Uncharacterized lipoprotein SACOL0083 (#Q5HJR9.1) (unknown/uncharacterized protein) |  |

### 3 Immunohistochemical staining of human tissue with specific anti-microbial antibodies

Frozen section slides of human tissues and the formalin-fixed paraffin-embedded mouse tissues were purchased commercially (https://www.biomax.us/index.php) and used directly for immunohistochemical staining according to the manufacturer’s instruction. The primary antibodies against EF-Tu of *Acinetobacter baumannii*, RPOB from *E.coli* and *hsp65* of *Mycobacterium tuberculosis* were diluted at 1:200, and applied to the frozen section tissue slides. The primary antibodies were incubated with the tissue slides for 1 hour at room temperature. The slides were washed 3 times with PBST and the secondary antibodies against the rabbit or the mouse IgG conjugated with horseradish peroxidase (HRP) were applied to the tissue slides. The signals were developed by using DAB enhanced detection system kit from Pierce (Fisher Scientific).

### 4 Sandwich enzyme-linked immunosorbent assay (ELISA assay)

Once the microbial protein identities were known, commercial antibodies were sought, and there were commercial antibodies against various proteins of different species. Depending upon the homology between the amino acid sequences, cross-species antibody may or may not bind to the same protein of another species. Based on the homology of amino acid sequences to human proteins shown in Table 1, a list of commercial antibodies were purchased (Table 2), and tested in the Sandwich ELISA assays.

**Table 2:** List of commercial antibodies from various species used in immunohistochemical staining and validation by sandwich ELISA assays.

Cross-species reactivity of antibodies
| • Microbial protein | Antibody immunogen | React to human |
| --- | --- | --- |
| • RPOB (*S. aureus*) | *E. coli* | Yes |
| • EF-G (*S. aureus/Pseud*) | Human | Yes |
| • HSP65 (*MAP/TB*) | *M. Tuberculosis* | Yes |
| • ATP synthase (*S.aureus/TB*) | Human | Yes |
| • EF-Tu (*E.coli*) | *Acinetobacter* | Yes |

Sandwich ELISA assay is a modified version of ELISA assay described as the following: The 96-well plates (ELISA plates, Fisher Scientific) were coated with the specific antibodies in Table 2 in coating buffer (at the concentration of 1.0 ug/ml) at 4°C overnight (Biolegend protocol online, http://www.biolegend.com/media_assets/support_protocol/BioLegend_Sandwich_ELISA_protocol.pdf). The coated 96-well plates were washed three times with PBST and the microbial whole cell extracts were added to the wells at the concentration of 20 ug/ml at room temperature for 2 hours. The plates were washed three times with PBST buffer and the whole plasmas of 50 ul were added from the patients with CD and SS as well as the normal healthy controls. The plasmas were incubated in the plate at room temperature for 1 hour. The plates were washed three times with PBST and the secondary antibody (anti-human IgG conjugated with horseradish peroxidase HRP) was added for 1 hour. The color development with tetramethylbenzydine reagent (TMB) (Biolegend, San Diego, CA) and the acid stop buffer were used and the absorbance reading at 450 nm was performed using Molecular Device Versamax microplate reader (Molecular Device, Sunnyvale, CA).

The plasma samples from the patients with Crohn’s disease and Sjogren’s syndrome were collected from various parts of the world through the commercial testing service at PZM Diagnostics (Charleston, WV, www.crohnsmanagement.com) with patients’ consent requisition forms and signatures on file. The healthy controls were obtained from the local individual clinic with the patients’ consent.

## Results

### 1 Identification of microbial proteins reactive to human plasma of Crohn’s patients

During our initial testing for MAP from the blood of patients with CD, we found a significant percentage of patients positive for the presence of antibody within the blood against MAP whole cell extract (over 70%, data not shown). We wanted to determine which specific antigen(s) from the MAP whole cell extract were reactive to human antibody in the plasmas of the patients. We performed Western blot analysis using the MAP whole cell extracts (20 ug) on the SDS polyacrylamide gel electrophoresis, and human plasma as primary antibodies (1:50 dilution). Surprisingly, we found that there was one prominent protein from MAP and MAH strongly reactive to patient’s plasma (Figure 1A, M60). There were multiple bacterial proteins from *S. aureus* and *S. pseudintermedius* reactive to the plasma of the same patient (S75, S27, S55). There were patients only reactive to proteins from *S. aureus* or *E. Coli* but not both (Figure 1B, S135, E39, E42).

We used the same patient’s plasma and the bacterial extracts to perform the immunoprecipitation and mass spectrometry to determine the identities of these bacterial/mycobacterial proteins reactive to patients’ plasmas. Nanospray LC mass Spectrometry and protein identification were performed at the Poochon Scientific LLC, Frederick, MD, and a list of potential bacterial/mycobacterial candidates from each immunoprecipitation sample was provided to us after the database search, and only the proteins with identical molecular weights to those on the Western blot were considered for further validation (Table 1). There were seven microbial proteins identified from the four microbes: S135 represents a bacterial protein from *S. aureus*, a strain of bacterium isolated from the blood of one patient with Crohn’s disease, and confirmed by partial 16S rDNA and whole genome sequencing by Illumina Miseq service (data not shown). Using a plasma from a 20 year old female with colonic Crohn’s disease as a primary antibody for Western blot analysis, S135 is demonstrated (Figure 1B). Immunoprecipitation using the whole cell extracts from the cultured *S. aureus* and the plasma from the same young female patient and mass spectrometry of the precipitated bacterial proteins showed the S135 to be DNA-directed RNA polymerase subunit B (RPOB, Genbank #ANR83811). RPOB is a critical enzyme for bacterial gene transcription. It is also a clinically relevant target for rifampin, one of the most important drugs for tuberculosis. Mutation of the RPOB gene confers rifampin resistance in *E. coli, S. aureus* and *M. tuberculosis* (12–14).

S75 represents a bacterial protein from *S. pseudintermedius*, a commensal bacteria on the surface of a domestic dog with potential to be pathogenic in humans (15, 16). The strain of *S. pseudintermedius* was isolated from a dog with a skin ulcer (hot spot, moist dermatitis), and the isolate was confirmed by partial 16S rDNA and whole genomic sequencing by Illumina Miseq service (data not shown). Using Western blot analysis and immunoprecipitation of plasmas of two separate patients with CD as the primary antibodies and using mass spectrometry, S75 was determined to be elongation factor G (EF-G, Genbank # WP_014614745) from *S. aureus* (Figure 1C). EF-G is a part of the 30S ribosome present in all bacteria species and it may relate to tetracycline resistance (17, 18). The human homologue of the bacterial EF-G is present in the mitochondria (G elongation factor mitochondrial 1, GFM1), and it plays similar roles in human protein biosynthesis. Human GFM1 gene mutation, can cause neonatal mitochondrial hepatoencephalopathy (19). The homology between the bacterial protein and the human protein at the amino acid levels is 61% by BLASTP search (Table 1).

M60 is a protein from MAP. M60 was identified by using MAP and MAH (*Mycobacterium avium subspecies hominissuis*) whole cell extracts and a plasma sample from a patient with CD (Figure 1A). The strain of MAH was isolated from the blood of the same patient by the mycobacterial culture, partial 16S rDNA and whole genomic DNA sequencing as previously described (10). M60 was determined to be heat shock protein 65 (*hsp65,* Genbank # P9WPE6). The human homologue of the mycobacterial *hsp65* is the human *hsp60*, a molecular chaperone known to be an autoantigen in cancer patients and other autoimmune diseases (20, 21).

S55 was a bacterial protein seen in *S. aureus* and *S. pseudintermedius* by using the plasma of a patient with Crohn’s disease (Figure 1 A). S55 was determined to be ATP synthase alpha from *S. aureus*, and *M. tuberculosis* (Genbank #KZK18041). The human homologue of ATP synthase alpha is present in the mitochondria (ATP5a), and it plays an important role in ATP generation and energy biosynthesis.

E42 was a bacterial protein from *E. coli* identified by Western blot using a plasma from a patient with mixed CD, SS and neurological symptoms such as decreased mental status and alertness. Further identification showed the bacterial protein to be consistent with elongation factor Tu (EF-Tu) from *E.coli* (Genbank # AAA50993). EF-Tu is one of the most abundant proteins from *E.coli*, and there is a human homologue of the bacterial EF-Tu from *E.coli*, EF-Tu mitochondrial precursor (TUFM, eEF1A1). Human eEF1A1 was identified as an autoantigen in 66% patients with Felty syndrome (rheumatoid arthritis, splenomegaly and neutropenia) and contributed to destruction of neutrophils and neutropenia (22).

E39 was a protein from *E.coli* reactive to plasma of the same patient with mixed CD and SS by Western blot analysis. The protein was identified as outer membrane porin C (ompC, Genbank #AAA23728) from *E. coli* by immunoprecipitation and mass spectrometry using the same patient’s plasma as in Western blot. The bacterial protein omp C from *E.coli* has been previously used in the Prometheus inflammatory bowel disease (IBD) panel (San Diego, CA) (23). There is no human homologue of ompC from *E.coli*, although there is still a possibility of cross-reactivity of human antibody in the circulation against omp C from *E.coli* with human tissues. It is reported that a significant percentage of CD patients (up to 40%) showed anti-ompC antibody (IgA) in their circulation (24).

S27 was a smaller protein from the *S. pseudintermedius* reactive to the plasma of a young female patient with Crohn’s disease. The protein was determined to be the uncharacterized lipoprotein SACOL0083 (Genbank accession #Q5HJR9.1). There is no previous documentation of this protein.

### 2 Specific anti-microbial antibodies cross-react to normal human tissues

To determine if the specific antibodies against the microbial proteins can react to human tissue, and the localization of the target antigens in human tissue, immunohistochemical staining techniques are employed. Specific antibodies from various sources were obtained commercially (Table 2, www.linscottsdirectory.com, Biolegend, San Diego, CA, Bethyl lab, TX, Mybiosorces, San Diego). Specific polyclonal antibody against EF-Tu from *Acinetobacter baumannii* in the rabbit can directly interact with human thyroid tissue with strong nuclear signal (Figure 2 A-C, A-negative control, B and C at 100X and 200 X magnification). Interestingly, the human homologue of the EF-Tu from *E.coli* is a mitochondrial precursor, and TUFM (human) is located within the mitochondria (cytoplasmic fraction), instead of nuclei. The significance of this finding is unclear. Specific monoclonal antibody from the mouse against *hsp65* from *M. tuberculosis* can also directly interact with human normal thyroid tissue (Figure 2, D-F, D-negative control at100X, E and F at 200 X magnification), and the reactivity signals were seen within the cytoplasm of the normal thyroid follicular cells. Polyclonal RPOB antibody only reacts to the skeletal muscle in a specific manner in which the signals were fine granules or dot in the cytoplasmic membrane (Figure 2, G-I, G-negative control x100, H and I at 200 X magnification). No nuclear signals were seen. None of the antibodies were reactive to the salivary gland tissue or the nucleated blood cells.

**Figure 2:**
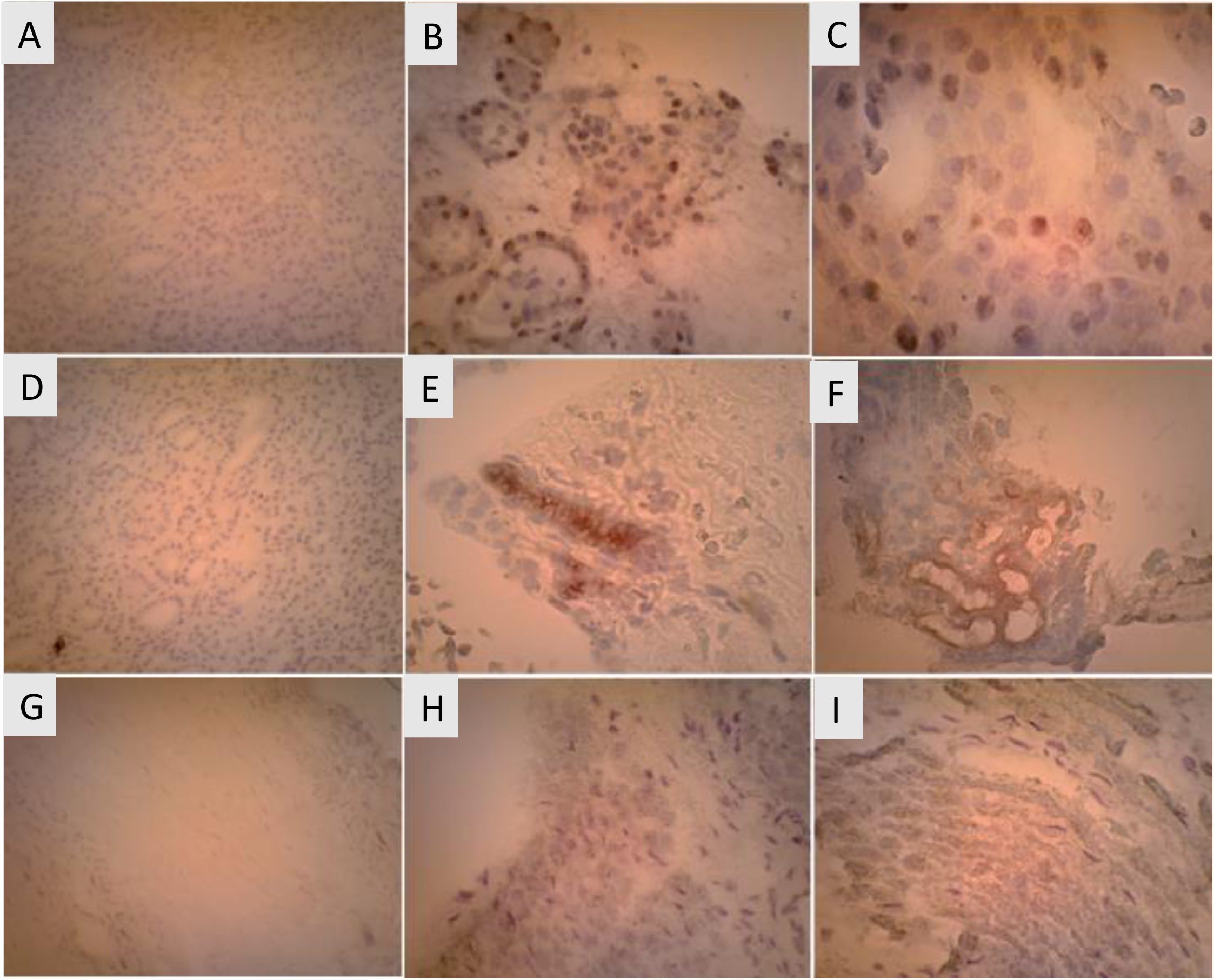
Cross-reactivity by immunohistochemical staining of the thyroid tissue and skeletal muscle. A-C: Polyclonal anti-EF-Tu antibody specific to *Acinetobacter baumannii* and thyroid tissue. A: negative control with no added antibody against EF-Tu at 100 X magnification with hematoxylin counterstain of the nuclei (blue); B: positive signals (brown, HRP with DAB system) were seen in the nuclei of the thyroid follicular cells at 200X magnification; C: positive signals were present in the nuclei of the thyroid cells at 200X magnification. D-E: Cross-reactivity of specific monoclonal antibody against *hsp65* from *M. tuberculosis* with the normal thyroid tissue. The signals were seen in the cytoplasm of normal thyroid follicular cells. D: negative control at 100 X, E and F: staining signals within the cytoplasm of the normal thyroid tissue at 200X magnifications. G-I: Cross-reactivity of specific polyclonal antibody against RPOB form *E.coli* with the skeletal muscle. The reactivity signals were seen as fine-granules and dots within the cytoplasm. G: negative control without antibody at 100X, H and I showed reactive signals within the cytoplasm and membranes in skeletal muscle at 200 X magnification.

### 3 Elevated levels of anti-microbial antibodies in patients with CD and SS

The specific polyclonal and monoclonal antibodies against the microbial proteins were purchased commercially (Table 2), and used for sandwich ELISA assays to determine the anti-microbial antibodies in the blood of the patients CD and SS. Clinical characteristics and the demographics of the CD patients and SS patients are listed in Table 3. In total, 44 plasma samples were available from the CD patients with 31 normal healthy controls, and 26 plasma samples from SS patients. The raw data (OD450 reading) of the plasma levels of anti-RPOB antibodies (IgG) in CD and the controls is shown in Figure 3. The cutoff value was the mean + 1 standard deviation (SD) of the normal control. The plasma levels of anti-RPOB IgG from the CD patients and the normal control were compared by using unpaired Student *t-test* and the presented with the boxplot as shown (Figure 3). In the CD patients, 41 patients’ plasma samples were positive for one or more markers (93%; all markers were positive in 14 patients, 4 markers were positive in 7 patients, 3 markers were positive in 8, 2 markers were positive in 8 patients, 1 marker was positive in 4 patients, all markers were negative in 3), 14 patients were positive for all five markers (32%), and 3 patients were negative for all the five markers (7%). In contrast, there are 6 patients positive for one or more markers in 31 controls (19%) (all markers were positive in 0 patient, one or more markers were positive in 6 patients) (Figure 4). The levels of all five markers were significantly elevated in CD patients compared to the normal controls by unpaired *Student t-test* (p=0.031 for *hsp65* to p<0.0001 RPOB and EF-Tu) as shown in the boxplot (Figure 4).

**Figure 3:**
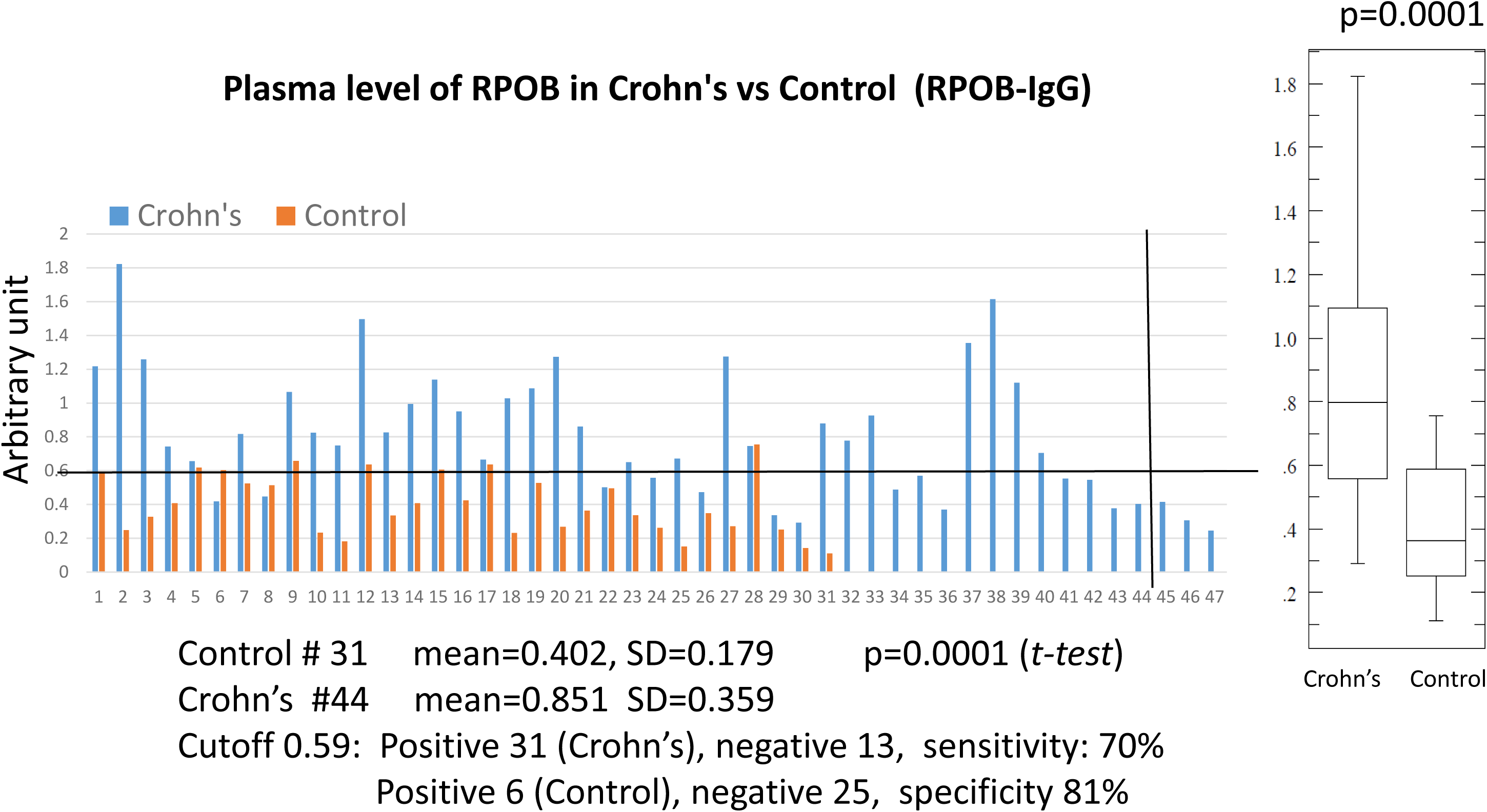
Plasma levels of anti-RPOB antibody in patients with Crohn’s and controls. The commercial anti-RPOB antibody was used to coat the 96-well plate and the whole *S. aureus* extracts were added to the wells with coated antibodies. Human plasmas from the Crohn’s patients and the controls were added to the wells and signals detected by HRP-conjugated anti-human IgG secondary antibody. The raw data was from OD450 readings from 44 CD patients and 31controls (left), and analyzed by using unpaired Student t-test with boxplot (right). The cutoff value was the mean + 1 standard deviation (SD).

**Figure 4:**
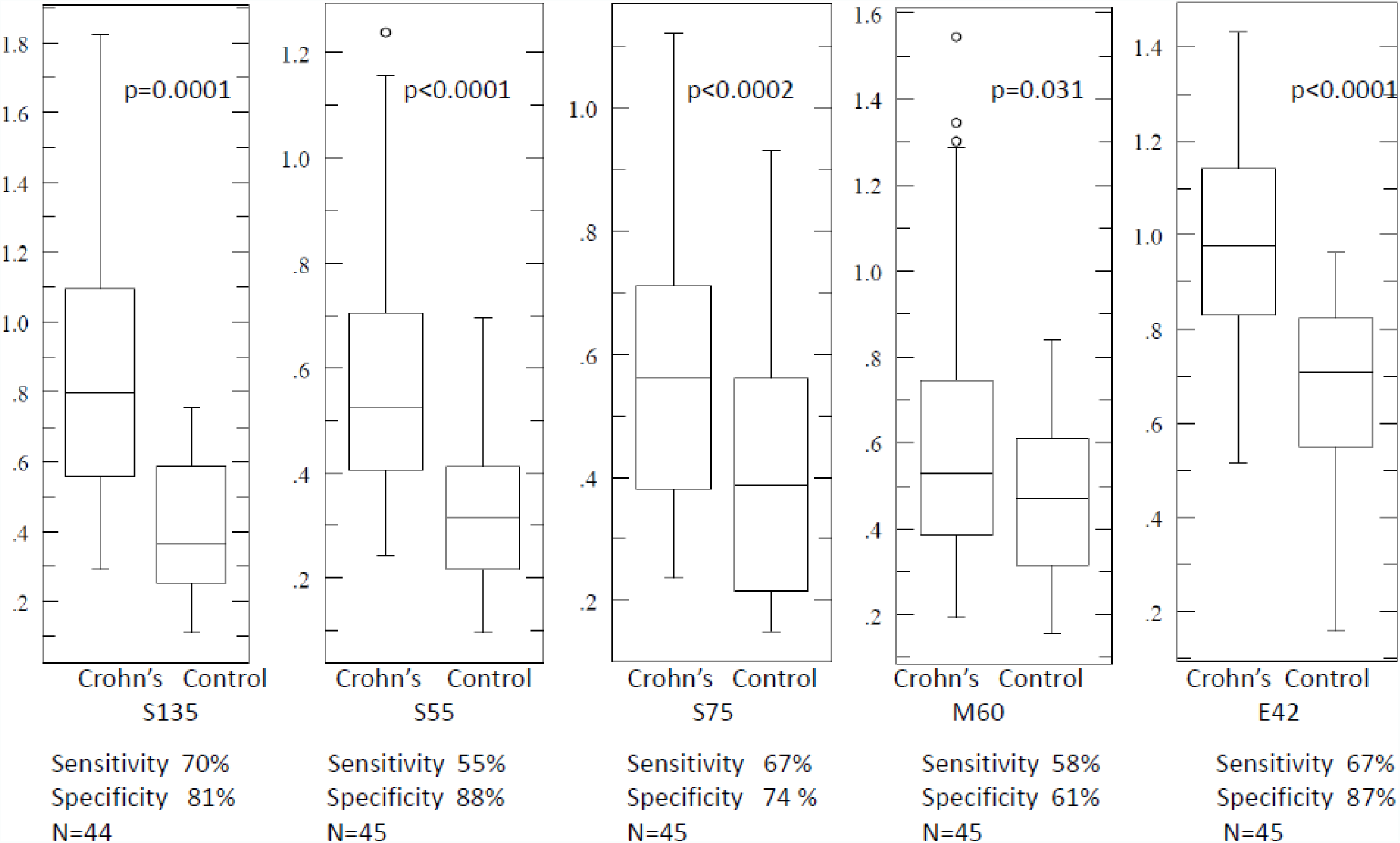
Plasma levels of anti-microbial antibodies against 5 bacterial/mycobacterial proteins in Crohn’s and the control. The statistical analyses were performed and the *p-values* were calculated by using unpaired Student t-test. Each antibody was performed and analyzed separately. Positive and negative values were defined as above or below the cutoff value (the mean + 1 SD). Sensitivity was expressed as the percentage of the patients with positive results versus the total CD patients, and specificity as percentage of negative results in normal controls versus the total controls.

**Table 3:** Clinical characteristics of Crohn’s patients and Sjogren’s patients in the study.

|  | Crohn's Disease | Sjogren's Syndrome |
| --- | --- | --- |
| Male/female | 30/14 (68/32%) | 5/21 (19/81%) |
| Median age at testing | 34 (5-71) | 58 (39-71) |
| Median duration of disease at testing | 7 (1-40) | Unknown |
| Endoscopy/histology | 44 (100%) | Unknown |
| Medication at testing |  | Unknown |
| Mesalamine–sulfasalazine | 2 |  |
| Corticosteroids | 11 |  |
| Immunosuppresants | 10 |  |
| Anti-TNF | 10 |  |
| Antibiotics (%) 0 (0) 9 (16.9) 0.033 | 14 |  |
| Methotrexate | 2 |  |
| Other | 7 |  |

Among the five serological markers, *hsp65* from *Mycobacterium* appears less differentiating with overlapping mean data points, and it is difficult to separate the patients versus the normal controls with large overlapping results and the borderline range, although the difference of the levels in patients and controls was significant by the unpaired *Student t-test* (p=0.031). All other four markers were easily separable between the CD patients and the normal controls with small overlapping ranges (Figure 4). The cutoff value in each marker was determined by the mean + 1 standard deviation of the control, and each cutoff value was determined separately based on the property of each antibody used (Figure 4). We did not correlate the disease severity, demographics, and medications with specific antibody levels, and we did not test if these serologic markers are useful for monitoring the treatment effect or efficacy.

In total, 26 patients’ plasmas from patients with SS were available for analysis, and 22 patients were positive for one or more markers (85%; 7 patients were positive for all markers, 5 patients were positive for 4 markers,5 patients were positive for 3 markers, 4 patients were positive for 2 markers, 1 was positive for 1 markers), and 4 patients were negative for all markers (17%) (Figure 5). All five markers showed statistically significant differences in the SS patients versus the controls by the unpaired *Student t-tests* (Figure 5).

**Figure 5:**
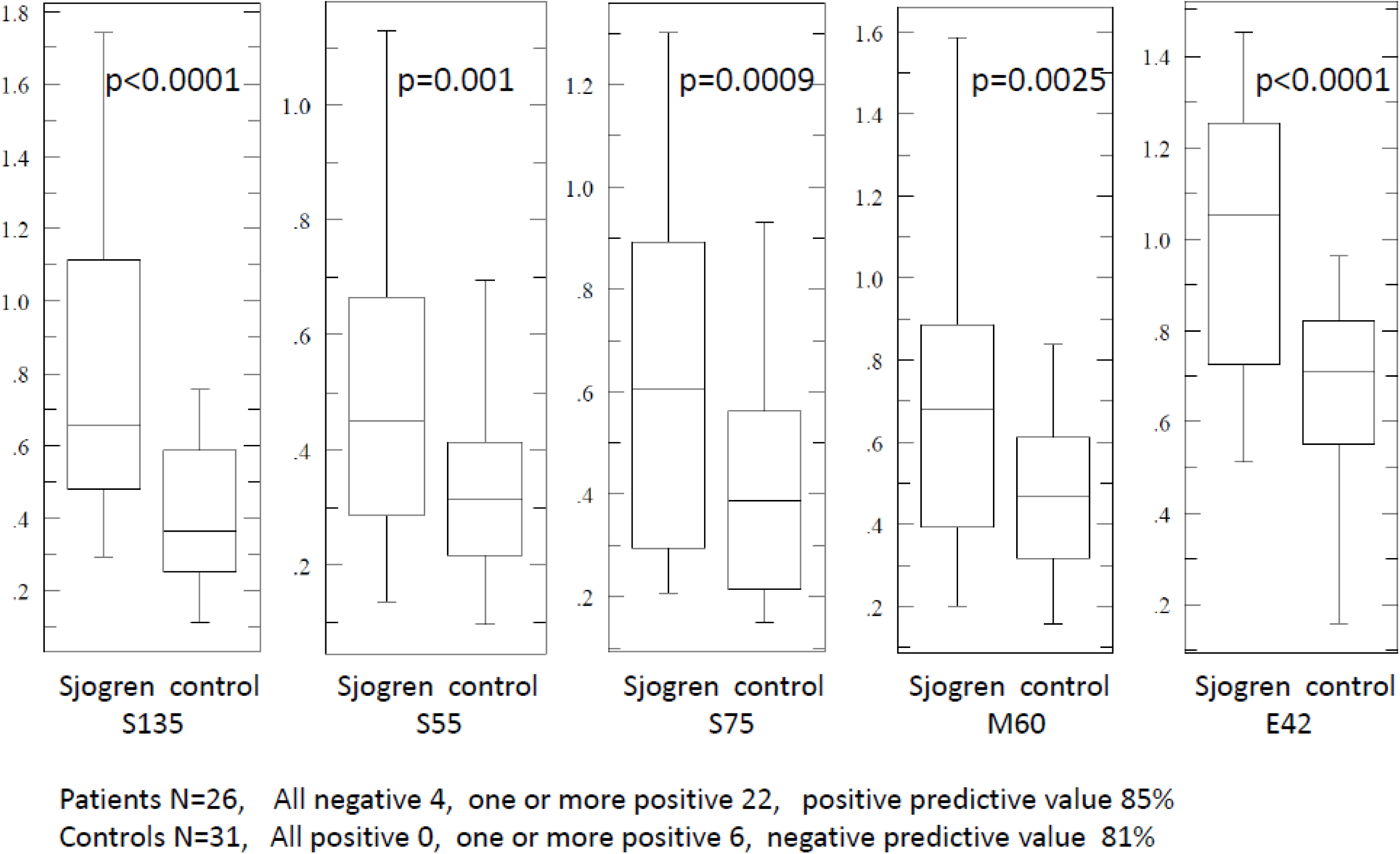
Plasma levels of anti-microbial antibodies against 5 bacterial/mycobacterial proteins in Sjogren’s patients and the control. The statistical analyses were performed and the *p-values* were calculated by using unpaired Student t-test. Each antibody was performed and analyzed separately. Positive and negative values were defined as above or below the cutoff value (the mean + 1 SD). Positive predictive value was expressed as the percentage of the patients with positive results versus the total SS patients, and negative predictive value as percentage of negative results in normal controls versus the total normal controls.

## Discussion

The patient population of all autoimmune diseases is increasing and the underlying causes of all these autoimmune diseases are unknown. Much of the effort is directed to discover the autoantibodies, and the treatment strategy is directed to suppress the immune system through various mechanisms. We have discovered a panel of antibodies against the microbial proteins and these microbes commonly cover the surface of the human body or in the case of mycobacteria, are widely present in the environment such as water and soil. *S. pseudintermedius* is a commensal bacterium on the surface of domestic dogs, and there are millions of families with domestic pets in the US.

There is a symbiotic relationship between the common microbes and the human hosts, and whether these common microbes become pathogenic to the host or not is likely determined by the genetic susceptibility of the individual. When there is a specific genetic variant of certain gene (s) leading to abnormal gene function, the individual is reactive with his/her own microbes within or on the surface of the body. In classic immunology, the development of specific circulating antibody against specific microbes is likely a result of the failed innate immunity network which consists of non-specific defense system such as body barriers including surface skin and mucosa lining through body cavities, and the blood defense system including neutrophils, macrophages, mast cells, eosinophils, and complement (1) (Figure 6). Specific cell wall proteins of bacteria, such as protein A from *S. aureus*, Protein G from Group G *Streptococcus*, Protein L from *Clostridium (Peptostreptococcus)*, and Protein D from *Haemophilus influenza/E. coli*, can bind to the immunoglobulins non-specifically with significantly high affinity, and these bacteria can be cleared rapidly through opsonization (25–29). The development of specific antibody against specific microbial proteins indicates the failure of clearance of the microbes from the circulation by the innate immune system network, and this failure to clear the microbes may directly reflect the genetic susceptibility to CD (30–33), SS and other autoimmune diseases. Defective functional activity of the mannose-binding lectin due to polymorphisms of mannose-binding lectin gene appears to correlate with the elevated levels of circulating anti-*Saccharomyces cerevisiae* (*S. cerevisiae*) antibody in Crohn’s patients (34, 35). *S. cerevisiae* has been demonstrated to be beneficial in the normal human gut microbiome (commensal) through metagenomics sequencing ^35^. It appears that anti-*S. cerevisiae* antibodies occurs in only certain Crohn’s patients with various sensitivity and specificity (36). Commensal intestinal bacteria, *Bacteroides fragilis*, can communicate with mucosal dendritic cells through noncanonical autophagy pathway in ATG16L1 and NOD2 dependent manner to induce regulatory T-cells (Tregs) in suppression of mucosal inflammation (31). Both ATG16L1 and NOD2 gene mutations/variations are major risk factors for CD (37). Whether these patients carry elevated anti-microbial antibodies, especially anti-*E.coli* antibodies or anti-*Bacteroides* antibodies in their circulation remains to be investigated, although CD patients with NOD2 gene variants are associated with elevated levels of anti-microbial antibodies especially anti-CBir-1 (38, 39). The molecular mechanism by which the circulating antibodies are generated within the host against the commensal microbes such as *S. aureus, E.coli* and *Mycobacterium* and the host genetic variations remains to be further elucidated. Recent studies indicate Crohn’s disease is a syndrome of dysbiosis, ie, imbalance of microbes in the gut. Dysbiosis may reflect decreased diversity of the gut microbes and an increase of particular one phylum of the gut microbes or a decrease of another (40–42). Regardless, dysbiosis reflects the growth status of the microbes inside the gut but outside the circulation, and the presence of specific antibodies against the microbial proteins in the blood indicates an invasion or penetration of the gut mucosal barrier into the blood circulation, leading to transient or persistent bacteremia.

**Figure 6:**
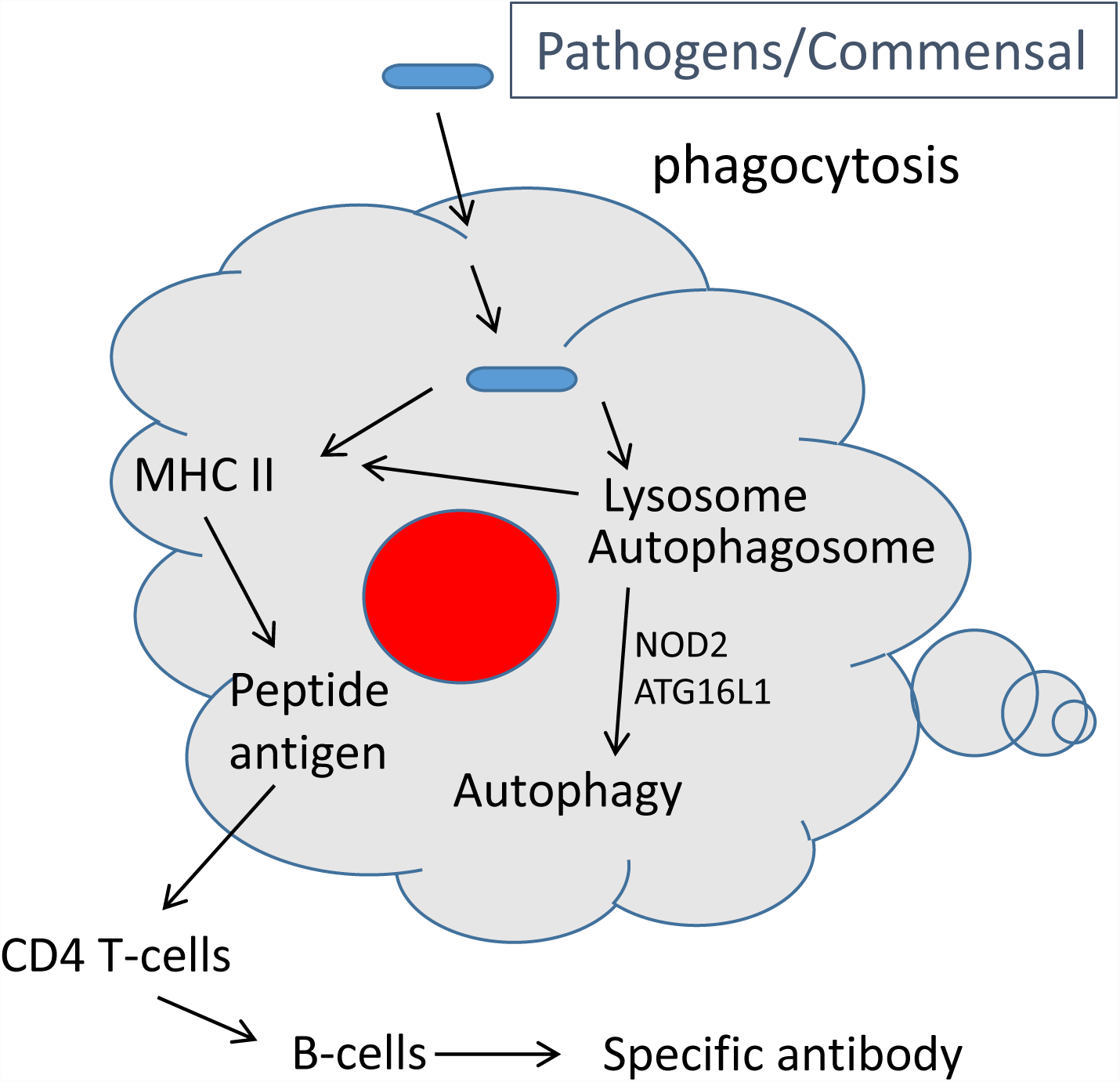
Schematic representation of innate immunity and adaptive immunity

We have discovered 7 proteins from 4 different types of microbes, *S. aureus, S. pseudintermedius, MAP*, and *E. coli*. *S. aureus* and *E. coli* are the most abundant bacteria on the body surface and in the gut. *S. pseudintermedius* is one of the most abundant bacteria in domestic dogs. *MAP* is present in the environment such as soil, water and dairy products. These microbial proteins are also highly homologous to the human counterparts (homologues), and the antibodies against the microbial proteins can directly interact with the human tissues. Our immunohistochemical staining results demonstrated that the specific monoclonal or polyclonal antibodies raised against the microbial proteins can cross-react to the human tissues. It is also important to note that there is significantly high homology of the amino acid sequences of these microbial proteins between the microbes such as *S. aureus* and *E. coli*, and the antibodies against *S. aureus* can also react to the same protein antigen from the *E. coli*. Cross-reactivity or molecular mimicry appears to be the key in understanding the mechanism of the underlying diseases caused by these common microbes. In diagnosis of Lyme disease, p58 on the Western blot positive for *Borrelia burgdorferi,* GroEL, is the same homologue as *hsp65* in *M. tuberculosis* and *E.coli* (43). EF-Tu from *B. burgdorferi* was found to be immunogenic during Lyme disease (44). It is plausible that the antibodies reacting to the extracts of *B. burgdorferi* in diagnosis of Lyme disease can result from the commensal bacteria such as *E. coli* or *S. aureus* within the body, instead of *B. burgdorferi* from the tick bite. Furthermore, the microbial proteins identified above are critical for survival of the microbes except for the unknown lipoprotein (S27) from the *S. aureus*. The elongation factors (EF-G and EF-Tu) are parts of the bacterial ribosomes targeted by some antibiotic families, and there is a remote possibility that the presence of the anti-microbial antibodies can result from the widespread antibiotics use to kill the bacteria, and the studies to understand the relationship of the anti-microbial antibodies with antibiotic usage as well as antibiotic resistance will be important. The homologues of the bacterial EF-G, EF-Tu, and ATP5a are all located within the mitochondria, and whether the anti-microbial antibodies cause mitochondrial dysfunction in a variety of human diseases such as chronic fatigue syndrome, metabolic syndrome, and neurodegenerative disorders will be of great interest.

In a clinical setting, testing anti-microbial antibodies can be used for confirmation of clinical diagnosis. Essentially, the anti-microbial antibody levels will indicate the failure of the innate immune system in the patient (failure to clear the microbes in blood). The presence of the antimicrobial antibodies potentially leads to the human tissue damage and the clinical diseases. A patient with clinical signs and symptoms of autoimmune diseases such as CD, SS, or other types of autoimmune diseases can be tested to see if there is/are elevated level(s) of one or more specific antibody (antibodies). The presence of the specific antibodies against the microbial proteins within the circulation of a patient suggests an entirely different mechanism of pathogenesis of autoimmune diseases.

Ultimately, autoimmune diseases are likely triggered by the infectious agents in genetically susceptible individual, and the infectious agents are commonly present on the surface of human body, or in the gut or in the environment. The discovery of the antibody panel against the microbial proteins in CD and SS may provide a new direction of research for understanding the autoimmune diseases, leading to new therapeutic strategy for a large population of patients with autoimmune diseases.

## References

1. Kumar, V., Abbas, AK., Aster, JC. 2015. Robbins and Cotran Pathologic Basis of Disease - 9th edition. Elsevier,

2. Tomkovich, S., and C. Jobin. 2016. Microbiota and host immune responses: a love-hate relationship. Immunology 147:1–10.

3. Grigg, J.B., and G.F. Sonnenberg. 2017. Host-Microbiota Interactions Shape Local and Systemic Inflammatory Diseases. J Immunol 198:564–571.

4. Valkenburg, H., Goslings, WRO, Bots, AW, de Moor, CE, Lorrier, JC. 1963. Attack Rates of Streptococcal Pharyngitis, Rheumatic Fever and Glomerulonephritis in the General Population — The Epidemiology of Streptococcal Pharvngitis in One Village during a Two-Year Period. New England Journal of Medicine 268:694–701.

5. Dale, J.B., and E.H. Beachey. 1985. Epitopes of streptococcal M proteins shared with cardiac myosin. J Exp Med 162:583–591.

6. Malkiel, S., L. Liao, M.W. Cunningham, and B. Diamond. 2000. T-Cell-dependent antibody response to the dominant epitope of streptococcal polysaccharide, N-acetyl-glucosamine, is cross-reactive with cardiac myosin. Infect Immun 68:5803–5808.

7. Quinn, A., K. Ward, V.A. Fischetti, M. Hemric, and M.W. Cunningham. 1998. Immunological relationship between the class I epitope of streptococcal M protein and myosin. Infect Immun 66:4418–4424.

8. Kees-Folts, D., A.B. Abt, R.E. Domen, and A.S. Freiberg. 2002. Renal failure after anti-D globulin treatment of idiopathic thrombocytopenic purpura. Pediatr Nephrol 17:91–96.

9. Sambrook, J., Fritsch, E.F., Maniatis, T. 1989. Molecular Cloning: A Laboratory Manual. Cold Spring Harbor Laboratory Press, New York.

10. Zhang, P., L.M. Minardi, J.T. Kuenstner, and R. Kruzelock. 2015. Mycobacterium avium subspecies hominissuis in Crohn’s disease: a case report. Gastroenterol Rep (Oxf) 1:1–4.

11. Zhang, P., L.M. Minardi, J.T. Kuenstner, S. Zekan, and R. Kruzelock. 2016. Extracellular Components in Culture Media of Mycobacterium Avium Subspecies and Staphylococci with Implications for Clinical Microbiology and Blood Culture. American Journal of Infectious Diseases and Microbiology 4:112–117.

12. Mokrousov, I., T. Otten, B. Vyshnevskiy, and O. Narvskaya. 2003. Allele-specific rpoB PCR assays for detection of rifampin-resistant Mycobacterium tuberculosis in sputum smears. Antimicrob Agents Chemother 47:2231–2235.

13. Telenti, A., P. Imboden, F. Marchesi, D. Lowrie, S. Cole, M.J. Colston, L. Matter, K. Schopfer, and T. Bodmer. 1993. Detection of rifampicin-resistance mutations in Mycobacterium tuberculosis. Lancet 341:647–650.

14. Wichelhaus, T.A., V. Schafer, V. Brade, and B. Boddinghaus. 1999. Molecular characterization of rpoB mutations conferring cross-resistance to rifamycins on methicillin-resistant Staphylococcus aureus. Antimicrob Agents Chemother 43:2813–2816.

15. Pompilio, A., S. De Nicola, V. Crocetta, S. Guarnieri, V. Savini, E. Carretto, and G. Di Bonaventura. 2015. New insights in Staphylococcus pseudintermedius pathogenicity: antibiotic-resistant biofilm formation by a human wound-associated strain. BMC Microbiol 15:109.

16. Somayaji, R., M.A. Priyantha, J.E. Rubin, and D. Church. 2016. Human infections due to Staphylococcus pseudintermedius, an emerging zoonosis of canine origin: report of 24 cases. Diagn Microbiol Infect Dis 85:471–476.

17. Burdett, V. 1991. Purification and characterization of Tet(M), a protein that renders ribosomes resistant to tetracycline. J Biol Chem 266:2872–2877.

18. Burdett, V. 1996. Tet(M)-promoted release of tetracycline from ribosomes is GTP dependent. J Bacteriol 178:3246–3251.

19. Ravn, K., B. Schonewolf-Greulich, R.M. Hansen, A.H. Bohr, M. Duno, F. Wibrand, and E. Ostergaard. 2015. Neonatal mitochondrial hepatoencephalopathy caused by novel GFM1 mutations. Mol Genet Metab Rep 3:5–10.

20. Kiessling, R., A. Gronberg, J. Ivanyi, K. Soderstrom, M. Ferm, S. Kleinau, E. Nilsson, and L. Klareskog. 1991. Role of hsp60 during autoimmune and bacterial inflammation. Immunol Rev 121:91–111.

21. Campanella, C., A. Marino Gammazza, L. Mularoni, F. Cappello, G. Zummo, and V. Di Felice. 2009. A comparative analysis of the products of GROEL-1 gene from Chlamydia trachomatis serovar D and the HSP60 var1 transcript from Homo sapiens suggests a possible autoimmune response. Int J Immunogenet 36:73–78.

22. Ditzel, H.J., Y. Masaki, H. Nielsen, L. Farnaes, and D.R. Burton. 2000. Cloning and expression of a novel human antibody-antigen pair associated with Felty’s syndrome. Proc Natl Acad Sci U S A 97:9234–9239.

23. Elkadri, A.A., J.M. Stempak, T.D. Walters, S. Lal, A.M. Griffiths, A.H. Steinhart, and M.S. Silverberg. 2013. Serum antibodies associated with complex inflammatory bowel disease. Inflamm Bowel Dis 19:1499–1505.

24. Dubinsky, M.C., Y.C. Lin, D. Dutridge, Y. Picornell, C.J. Landers, S. Farrior, I. Wrobel, A. Quiros, E.A. Vasiliauskas, B. Grill, D. Israel, R. Bahar, D. Christie, G. Wahbeh, G. Silber, S. Dallazadeh, P. Shah, D. Thomas, D. Kelts, R.M. Hershberg, C.O. Elson, S.R. Targan, K.D. Taylor, J.I. Rotter, and H. Yang. 2006. Serum immune responses predict rapid disease progression among children with Crohn’s disease: immune responses predict disease progression. Am J Gastroenterol 101:360–367.

25. Hjelm, H., K. Hjelm, and J. Sjoquist. 1972. Protein A from Staphylococcus aureus. Its isolation by affinity chromatography and its use as an immunosorbent for isolation of immunoglobulins. FEBS Lett 28:73–76.

26. Verwey, W.F. 1940. A Type-Specific Antigenic Protein Derived From The Staphylococcus. J Exp Med 71:635–644.

27. Akerstrom, B., T. Brodin, K. Reis, and L. Bjorck. 1985. Protein G: a powerful tool for binding and detection of monoclonal and polyclonal antibodies. J Immunol 135:2589–2592.

28. Akerstrom, B., and L. Bjorck. 1989. Protein L: an immunoglobulin light chain-binding bacterial protein. Characterization of binding and physicochemical properties. J Biol Chem 264:19740–19746.

29. Janson, H., L.O. Heden, and A. Forsgren. 1992. Protein D, the immunoglobulin D-binding protein of Haemophilus influenzae, is a lipoprotein. Infect Immun 60:1336–1342.

30. Jostins, L., S. Ripke, R.K. Weersma, R.H. Duerr, D.P. McGovern, K.Y. Hui, J.C. Lee, L.P. Schumm, Y. Sharma, C.A. Anderson, J. Essers, M. Mitrovic, K. Ning, I. Cleynen, E. Theatre, S.L. Spain, S. Raychaudhuri, P. Goyette, Z. Wei, C. Abraham, J.P. Achkar, T. Ahmad, L. Amininejad, A.N. Ananthakrishnan, V. Andersen, J.M. Andrews, L. Baidoo, T. Balschun, P.A. Bampton, A. Bitton, G. Boucher, S. Brand, C. Buning, A. Cohain, S. Cichon, M. D’Amato, D. De Jong, K.L. Devaney, M. Dubinsky, C. Edwards, D. Ellinghaus, L.R. Ferguson, D. Franchimont, K. Fransen, R. Gearry, M. Georges, C. Gieger, J. Glas, T. Haritunians, A. Hart, C. Hawkey, M. Hedl, X. Hu, T.H. Karlsen, L. Kupcinskas, S. Kugathasan, A. Latiano, D. Laukens, I.C. Lawrance, C.W. Lees, E. Louis, G. Mahy, J. Mansfield, A.R. Morgan, C. Mowat, W. Newman, O. Palmieri, C.Y. Ponsioen, U. Potocnik, N.J. Prescott, M. Regueiro, J.I. Rotter, R.K. Russell, J.D. Sanderson, M. Sans, J. Satsangi, S. Schreiber, L.A. Simms, J. Sventoraityte, S.R. Targan, K.D. Taylor, M. Tremelling, H.W. Verspaget, M. De Vos, C. Wijmenga, D.C. Wilson, J. Winkelmann, R.J. Xavier, S. Zeissig, B. Zhang, C.K. Zhang, H. Zhao, M.S. Silverberg, V. Annese, H. Hakonarson, S.R. Brant, G. Radford-Smith, C.G. Mathew, J.D. Rioux, E.E. Schadt, M.J. Daly, A. Franke, M. Parkes, S. Vermeire, J.C. Barrett, and J.H. Cho. 2012. Host-microbe interactions have shaped the genetic architecture of inflammatory bowel disease. Nature 491:119–124.

31. Chu, H., A. Khosravi, I.P. Kusumawardhani, A.H. Kwon, A.C. Vasconcelos, L.D. Cunha, A.E. Mayer, Y. Shen, W.L. Wu, A. Kambal, S.R. Targan, R.J. Xavier, P.B. Ernst, D.R. Green, D.P. McGovern, H.W. Virgin, and S.K. Mazmanian. 2016. Gene-microbiota interactions contribute to the pathogenesis of inflammatory bowel disease. Science 352:1116–1120.

32. Lee, J.C., D. Biasci, R. Roberts, R.B. Gearry, J.C. Mansfield, T. Ahmad, N.J. Prescott, J. Satsangi, D.C. Wilson, L. Jostins, C.A. Anderson, J.A. Traherne, P.A. Lyons, M. Parkes, and K.G. Smith. 2017. Genome-wide association study identifies distinct genetic contributions to prognosis and susceptibility in Crohn’s disease. Nat Genet 49:262–268.

33. Murthy, A., Y. Li, I. Peng, M. Reichelt, A.K. Katakam, R. Noubade, M. Roose-Girma, J. DeVoss, L. Diehl, R.R. Graham, and M. van Lookeren Campagne. 2014. A Crohn’s disease variant in Atg16l1 enhances its degradation by caspase 3. Nature 506:456–462.

34. Choteau, L., F. Vasseur, F. Lepretre, M. Figeac, C. Gower-Rousseau, L. Dubuquoy, D. Poulain, J.F. Colombel, B. Sendid, and S. Jawhara. 2016. Polymorphisms in the Mannose-Binding Lectin Gene are Associated with Defective Mannose-Binding Lectin Functional Activity in Crohn’s Disease Patients. Sci Rep 6:29636.

35. Hoffmann, C., S. Dollive, S. Grunberg, J. Chen, H. Li, G.D. Wu, J.D. Lewis, and F.D. Bushman. 2013. Archaea and fungi of the human gut microbiome: correlations with diet and bacterial residents. PLoS One 8:e66019.

36. Sendid, B., J.F. Colombel, P.M. Jacquinot, C. Faille, J. Fruit, A. Cortot, D. Lucidarme, D. Camus, and D. Poulain. 1996. Specific antibody response to oligomannosidic epitopes in Crohn’s disease. Clin Diagn Lab Immunol 3:219–226.

37. Travassos, L.H., L.A. Carneiro, M. Ramjeet, S. Hussey, Y.G. Kim, J.G. Magalhaes, L. Yuan, F. Soares, E. Chea, L. Le Bourhis, I.G. Boneca, A. Allaoui, N.L. Jones, G. Nunez, S.E. Girardin, and D.J. Philpott. 2010. Nod1 and Nod2 direct autophagy by recruiting ATG16L1 to the plasma membrane at the site of bacterial entry. Nat Immunol 11:55–62.

38. Devlin, S.M., H. Yang, A. Ippoliti, K.D. Taylor, C.J. Landers, X. Su, M.T. Abreu, K.A. Papadakis, E.A. Vasiliauskas, G.Y. Melmed, P.R. Fleshner, L. Mei, J.I. Rotter, and S.R. Targan. 2007. NOD2 variants and antibody response to microbial antigens in Crohn’s disease patients and their unaffected relatives. Gastroenterology 132:576–586.

39. Papadakis, K.A., H. Yang, A. Ippoliti, L. Mei, C.O. Elson, R.M. Hershberg, E.A. Vasiliauskas, P.R. Fleshner, M.T. Abreu, K. Taylor, C.J. Landers, J.I. Rotter, and S.R. Targan. 2007. Anti-flagellin (CBir1) phenotypic and genetic Crohn’s disease associations. Inflamm Bowel Dis 13:524–530.

40. Pascal, V., M. Pozuelo, N. Borruel, F. Casellas, D. Campos, A. Santiago, X. Martinez, E. Varela, G. Sarrabayrouse, K. Machiels, S. Vermeire, H. Sokol, F. Guarner, and C. Manichanh. 2017. A microbial signature for Crohn’s disease. Gut 66:813–822.

41. Schaubeck, M., T. Clavel, J. Calasan, I. Lagkouvardos, S.B. Haange, N. Jehmlich, M. Basic, A. Dupont, M. Hornef, M. von Bergen, A. Bleich, and D. Haller. 2016. Dysbiotic gut microbiota causes transmissible Crohn’s disease-like ileitis independent of failure in antimicrobial defence. Gut 65:225–237.

42. Mottawea, W., C.K. Chiang, M. Muhlbauer, A.E. Starr, J. Butcher, T. Abujamel, S.A. Deeke, A. Brandel, H. Zhou, S. Shokralla, M. Hajibabaei, R. Singleton, E.I. Benchimol, C. Jobin, D.R. Mack, D. Figeys, and A. Stintzi. 2016. Altered intestinal microbiota-host mitochondria crosstalk in new onset Crohn’s disease. Nat Commun 7:13419.

43. Carreiro, M.M., D.C. Laux, and D.R. Nelson. 1990. Characterization of the heat shock response and identification of heat shock protein antigens of Borrelia burgdorferi. Infect Immun 58:2186–2191.

44. Carrasco, S.E., Y. Yang, B. Troxell, X. Yang, U. Pal, and X.F. Yang. 2015. Borrelia burgdorferi elongation factor EF-Tu is an immunogenic protein during Lyme borreliosis. Emerg Microbes Infect 4:e54.

